# Electroencephalogram Data Collection for Student Engagement Analysis with Audio-Visual Content

**DOI:** 10.1101/2022.09.20.508656

**Authors:** Mansi Singh, Shreya Mishra, Jayantika Mehta, Anusha Bansal, Manoj Baranwal, Rahul Upadhyay, Vinay Kumar

**Author notes:** **Corresponding author**, Rahul Upadhyay.

## Abstract

Recognizing and monitoring students’ attention during learning is crucial to successful knowledge acquisition since it influences cognitive function. As a result, gaining a precise picture of a learner’s mental state may enable interactive learning systems to alter tutoring content, devise effective help tactics, and improve learning outcomes. In computer-based learning environments, keeping track of students’ mental states is vital. Investigating the feasibility of utilizing active learning in enhancing student engagement index when exposed to various visual stimuli is the genesis of this work. The research includes collecting EEG data from 20 participants (ten males, ten females) while resting and being subjected to various virtual infotainment/educational content. The EEG data were collected using the Allengers Neuro PLOT, a 40-channel wet electrode system. The work includes raw and pre-processed EEG data under quiescent and audio-visual continuous cues. The recorded data is accommodated by a sophisticated EEG data pre-processing pipeline and will be available to the research community for usage.

**Specifications Table:** 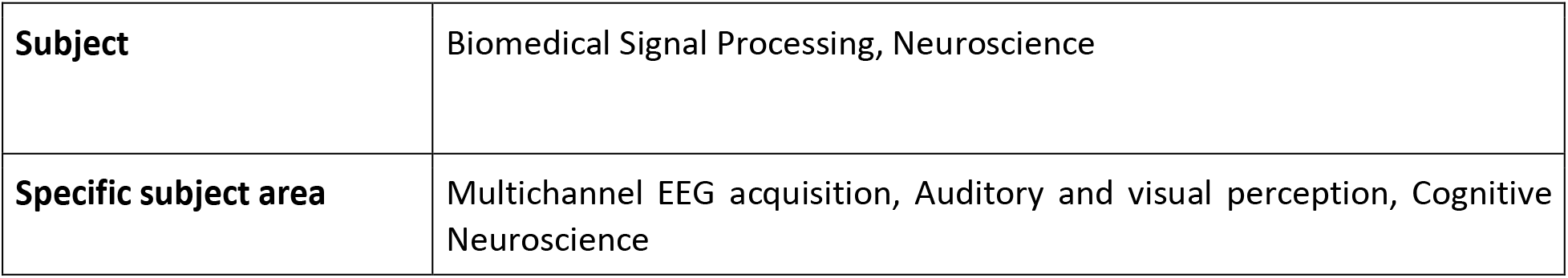

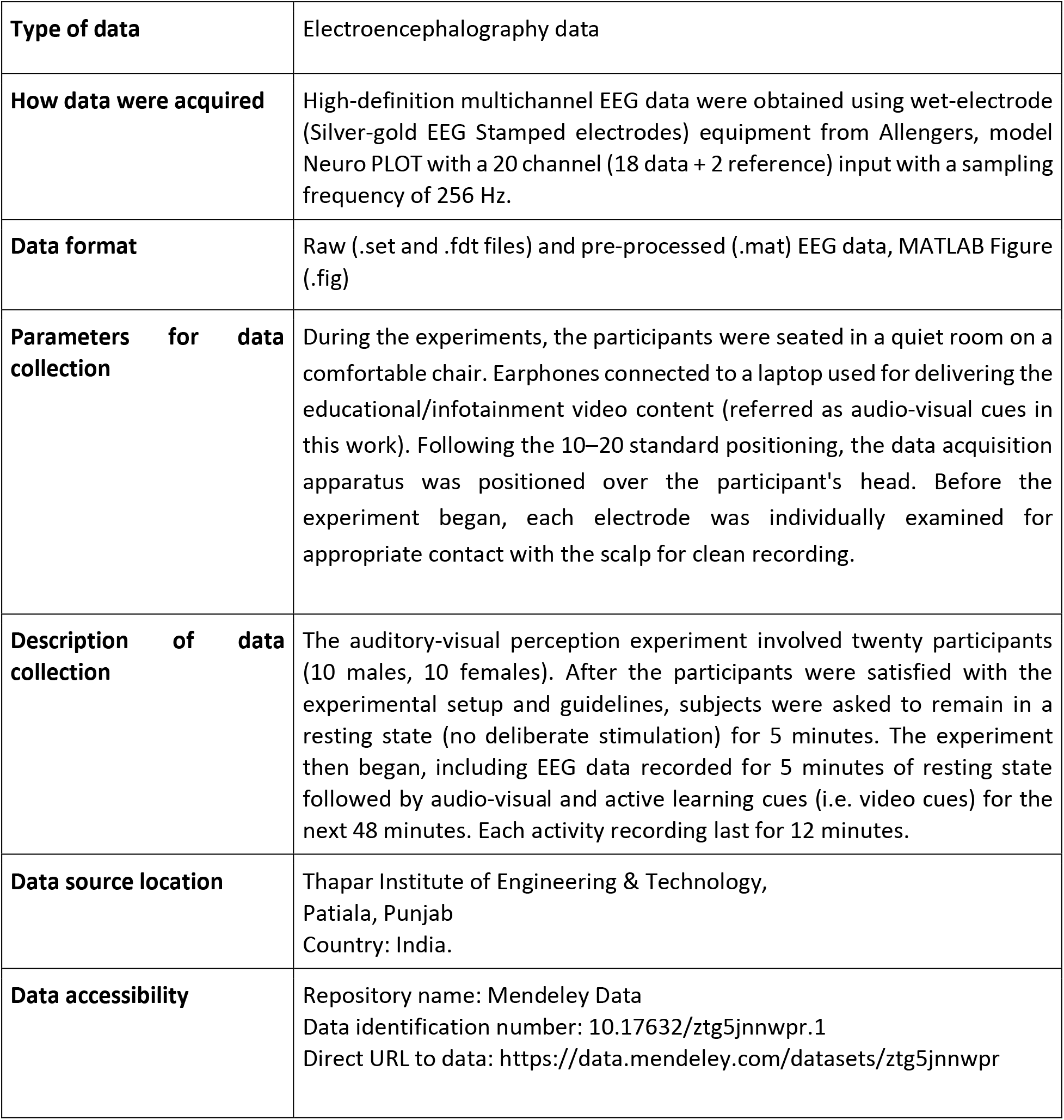

## Value of the Data

- In computer-based learning environments, it’s very important to keep track of students’/learners’ (referred as subjects) mental states. Because it affects subjects’ cognitive function, the capacity to notice and monitor their attention during the learning process is critical to successful knowledge acquisition (Khedher et al., 2019). Quantitative metrics in education may be used to find processes that improve learning efficiency, better align services with subjects’ requirements, or monitor critical task performance.
- Engagement may be the most important factor in making online learning a necessary element of higher education and an institution’s future. Student involvement promotes online course performance by increasing student contentment, increasing student drive to learn, reducing isolation, and increasing student satisfaction.
- For teachers who teach large online classes, the data and analysis can help to formulate tutoring content that is more interactive and engaging. Psychiatrists can utilise this information to analyse their patients’ brain activity. It can also be utilised as a self- or parent-report of academic performance, as well as the neurophysiological mechanisms underlying basic creative behaviours like idea development and appraisal.

## Data Description

- During the experiment, twenty volunteers (10 males, 10 females) took part in the experiment and EEG signals were recorded using an Allengers Neuro PLOT device with a 20 channel (18 data channels + 2 reference) with sampling frequency of 256 Hz. The eighteen channels include FP1, FP2, F3, F4, C3, C4, P3, P4, T3, T4, A1, A2, O1, O2, Fz, Cz, Pz, Oz as depicted in the Fig. 1. All participants were students, aged between 18-23 years and average age was 21 years. The raw and pre-processed data includes EEG data recorded for 5 minutes of resting-state followed by audio-visual stimuli-based data record of 48 minutes.
- For ease to post-processing and analysis, the pre-processed and filtered EEG data is provided in MATLAB’s ‘.mat’ file format, and raw data is included in .set and .fdt format. The EEG pre-processing pipeline was devised and used to process raw EEG activity i.e., free from artifacts and noise.

**Fig. 1:**
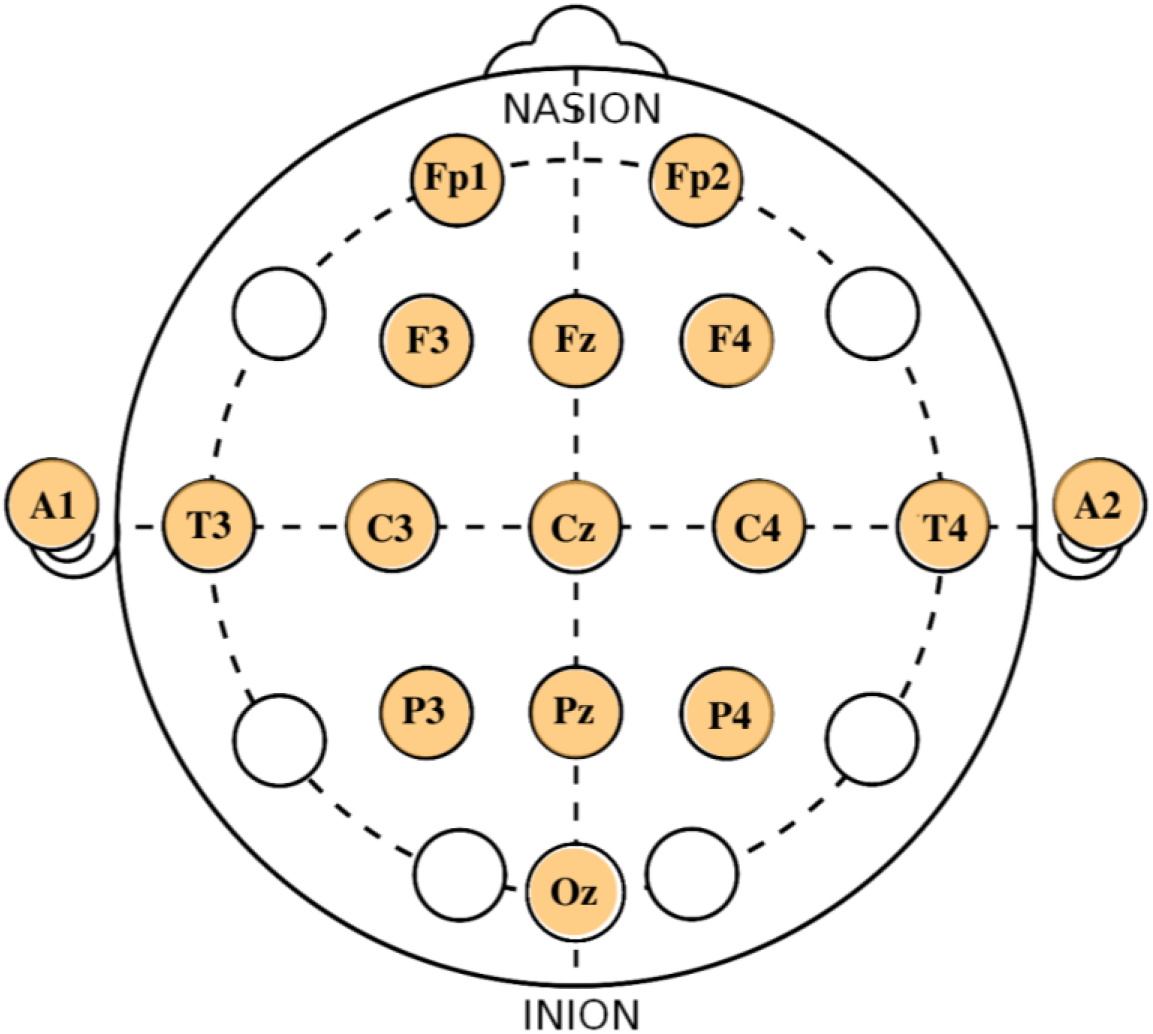
The electrode placement in Allengers NeuroPLOT (the electrodes utilised are highlighted in yellow colour using 10–20 electrode placement system)

## Experimental Design, Materials, and Methods

### Statement of Ethics

A voluntary, comprehensive informed consent was obtained/acquired from the participants. The Informed Consent form was approved by the Institutional Ethical Committee (IEC) of Thapar Institute of Engineering & Technology, Patiala and fulfilled the ethical guidelines outlined in the Declaration of Helsinki, 1964. There is no personally identifying information in the anonymised data.

### Participants

The electroencephalography data was recorded from 20 participants (10 male and 10 female). All participants were students of Thapar Institute of Engineering & Technology, Patiala and aged between 18-23 years.

### Risk and inconvenience

The following are the rare and minor physical risks associated with participating in this study:

a. possible skin irritation from the electrolyte paste or adhesive discs we use to place the electrodes on your skin;
b. discomfort from sitting still in one position for 53 minutes;
c. discomfort from electrolyte gel in subjects’ hair;
d. the possibility of receiving undetectable electrical signals from the equipment. One electrode (out of 18) carries a return current to the subject’s scalp, and electrical current is inherently limited to 50 microAmps, the chances of encountering electric discharge from the EEG apparatus are extremely minimal (lower thresholds of perceptible electric discharge are 1 milliAmp).

### Material and equipment

Allengers NeuroPLOT, a portable and commercially available EEG recording device, from Allengers medical systems ltd. was used to collect EEG activity. With 40 electrodes, this device records high-quality multichannel EEG data (includes EEG channels, Bipolar channels and reference channels). The device’s sampling rates are 256 Hz and 524 Hz. This equipment contains a 24 bit Analog to Digital Conversion (ADC), a full HD display of EEG waves at 1920 x 1080 pixels and can be used for analysing power spectra as well as coherence. The wet electrodes used are silver-gold EEG stamped disc electrodes from Spes Medica with Touch proof DIN 42802 connector. WEAVER and company’s Nuprep skin prep gel prepped the scalp and Ten20 conductive paste helped in pasting the disc electrodes on the scalp. The apparatus is linked to an exclusive software NeuroPLOT EEG installed on a desktop with an Intel Core i7 processor, 16 GB of RAM, and a 64-bit Windows operating system.

## Experiment

The study lasted 53 minutes in total, including the baseline as represented in Fig. 2. There were five steps to the experiment:

- ***Stage 1:* Baseline (B)**: There were no conscious stimuli used and subjects were asked to relax with minimum physical activity for 5-minutes. Following the rest period, the experiment continued, which comprised four 12-minutes of audio-visuals stimuli (video content) recordings corresponding to four different experiments as explained in stage 2-4.
- ***Stage 2:* Infotainment:** Broadcast content that was both entertaining and informative was played and EEG activity was recorded for 12 minutes.
- ***Stage 3:* Entertainment:** A video that held the subject’s attention and interest while also providing pleasure was played and EEG activity was recorded.
- ***Stage 4:* Active Learning:** The subject was handed a piece of paper and a pencil. Students were actively or experientially involved in the learning process. The subject was instructed to take notes as tutored in the video and simultaneously the EEG activity was recorded.
- ***Stage 5:* Educational Content:** The subject was asked to watch video content that was intended to educate them (related to course curriculum) and EEG activity was recorded. The educational video content was biased to the subjects for gaining maximum attention.

**Fig. 2:**
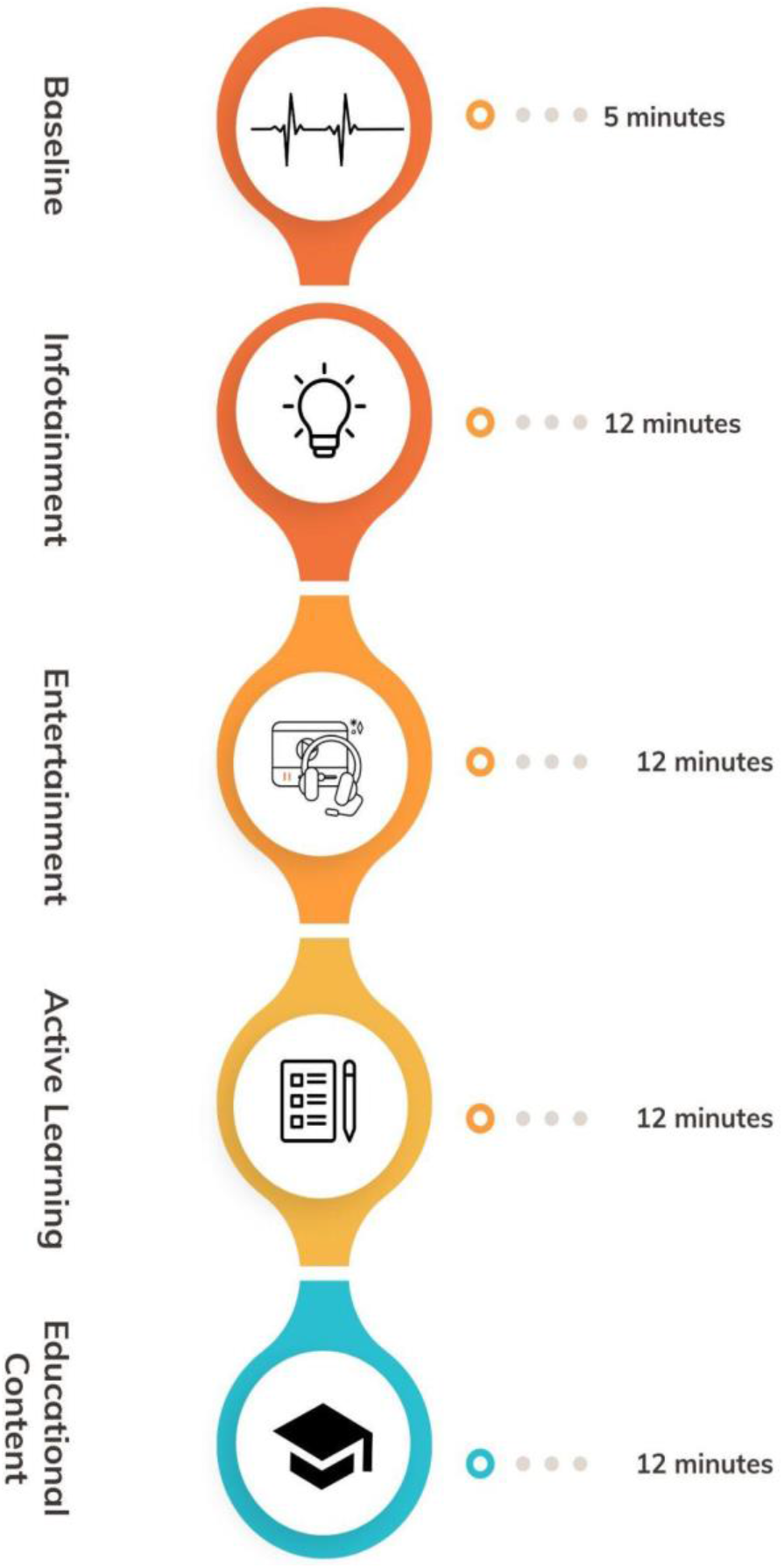
Experimental Design Protocol and timing diagram

## Experimental protocol

1. The complete experiment detail was clearly and precisely communicated to all participants. They were given instructions on how to conduct themselves during the experiment, such as avoiding unexpected movements, anxious thoughts, and talking.
2. All materials and equipment were organised and ready to use before the experiment began. The recording equipment will be tested to confirm that it is properly working.
3. The data recording apparatus was positioned above the participant’s head in the normal 10–20 position (Homan et al., 1987). Before the experiment began, each electrode was checked for appropriate contact and impedance matching to the scalp.
4. Once the subject was acquainted with the experimental setup and procedures, the trial began, with four audio-visual videos being shown and data was collected simultaneously.
5. Disturbances such as muscular activity, eye movements and blinks, physical movements, and external noise were removed from the EEG data and processed EEG activity is provided for analysis by the scientific community. The EEG data processing was accomplished in MATLAB software package. Fig. 3 shows the schematic of EEG data recording and processing methodology.

**Fig. 3:**
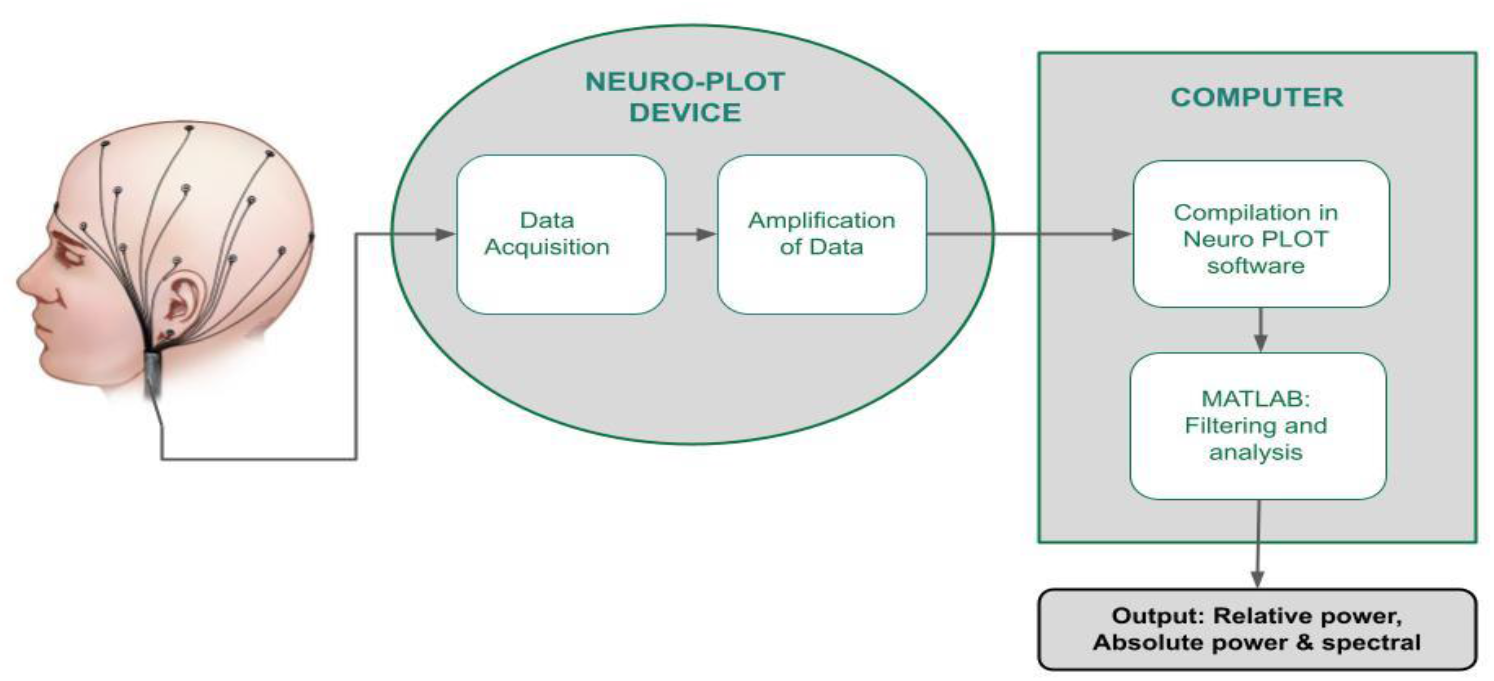
Schematic diagram of EEG data recording and processing

## Data pre-processing

In present work, Matlab R2019 (a) was used for offline processing and analysis of EEG data. The EEG data pre-processing was carried out using the Matlab plugin EEGLAB (Delorme and Makeig, 2004). The pre-processing pipeline is illustrated in Fig. 4. The raw data was imported as a .set file to EEGLAB using MATLAB. A bandpass filter of 4-45 Hz was used to filter the 18-channel EEG data. The goal is to improve the signal-to-noise ratio while lowering signal distortions. The location of EEG channels were imported in the EEGLAB and “automatic channel rejection” was used to objectively reject bad EEG channels and maintain channel quality control across participants. Following channel rejection, the removed channels were interpolated using the EEGLAB function “interpolate electrodes”. After interpolating the bad channels, the data was re-referenced to the common average. Further, an independent component analysis (ICA) was performed to generate topographical scalp-maps of the EEG source for individual components and decompose EEG into fundamental independent components for noise/artifact rejection. A logistic infomax technique with the natural gradient feature is an inbuilt ICA algorithm into EEGLAB (Sareen et al., 2020) which is used of ICA decomposition. Once independent components are isolated using infomax method, the EEGLAB plugin ADJUST was used to identify and remove independent components containing artefacts. ADJUST is an automatic algorithm that uses spatial and temporal features designed to recognise and remove eye blinks, eye movements and generic discontinuities (Mognon et al., 2011). Darbeliai is an EEGLAB plugin designed to allow batch processing of EEG data files in a graphical user interface (GUI). Darbeliai accepts EEG data files in a variety of formats (*.set, *.edf, *.cnt, and others) as input and outputs processed data to files in the EEGLAB data format (*.set), as opposed to EEGLAB itself, which requires users to interact with EEG datasets. Pictorial representation of the Absolute and Spectrum were be obtained.

**Fig. 4:**
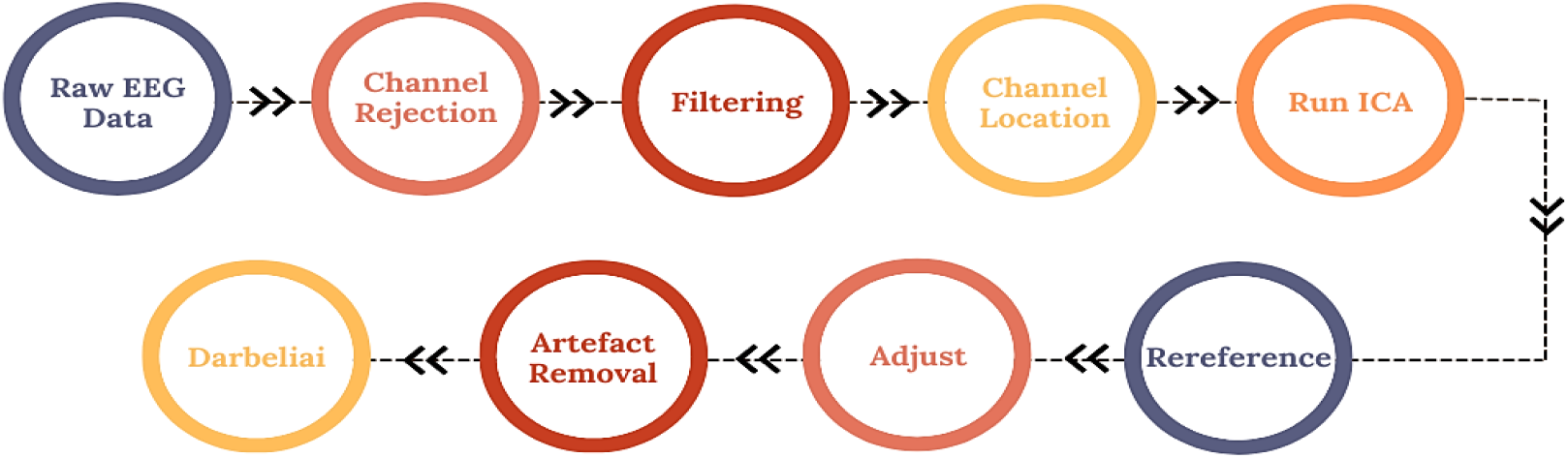
EEG data processing pipeline

## Observations

The pre-processed EEG activity is analysed to deduce significant pattern distinction in video stimulus engagement tasks. Fig. 5 and Fig. 6 represent the power spectrum and absolute power graphs obtained for entertainment, active Learning, infotainment and educational video content stimuli. It can be observed from Fig. 5(a) and Fig 5(b) that the subject exposed to the active learning and entertainment visual stimulus show significant EEG activity in the high frequency band (beta rhythm). Very negligible drop in EEG activity is witnessed during transition from low frequency rhythm i.e. delta and theta to high frequency rhythms. The high amplitude in the beta rhythm represents elevated levels of alertness in the subjects during active learning and entertainment visual stimulations. The high levels reflect active engagement of the subjects with the visual tasks. From Fig. 6(a) and Fig. 6(b), it is evident that for few channels we get significantly high value of absolute power compared to delta and theta bands. In Fig. 5(c) and Fig. 5(d), a noticeable drop in the EEG activity can be witnessed for high frequency rhythms for infotainment and education video content. Specifically, the drop is significant in few channels as evident from Fig. 5(d).

**Fig. 5:**
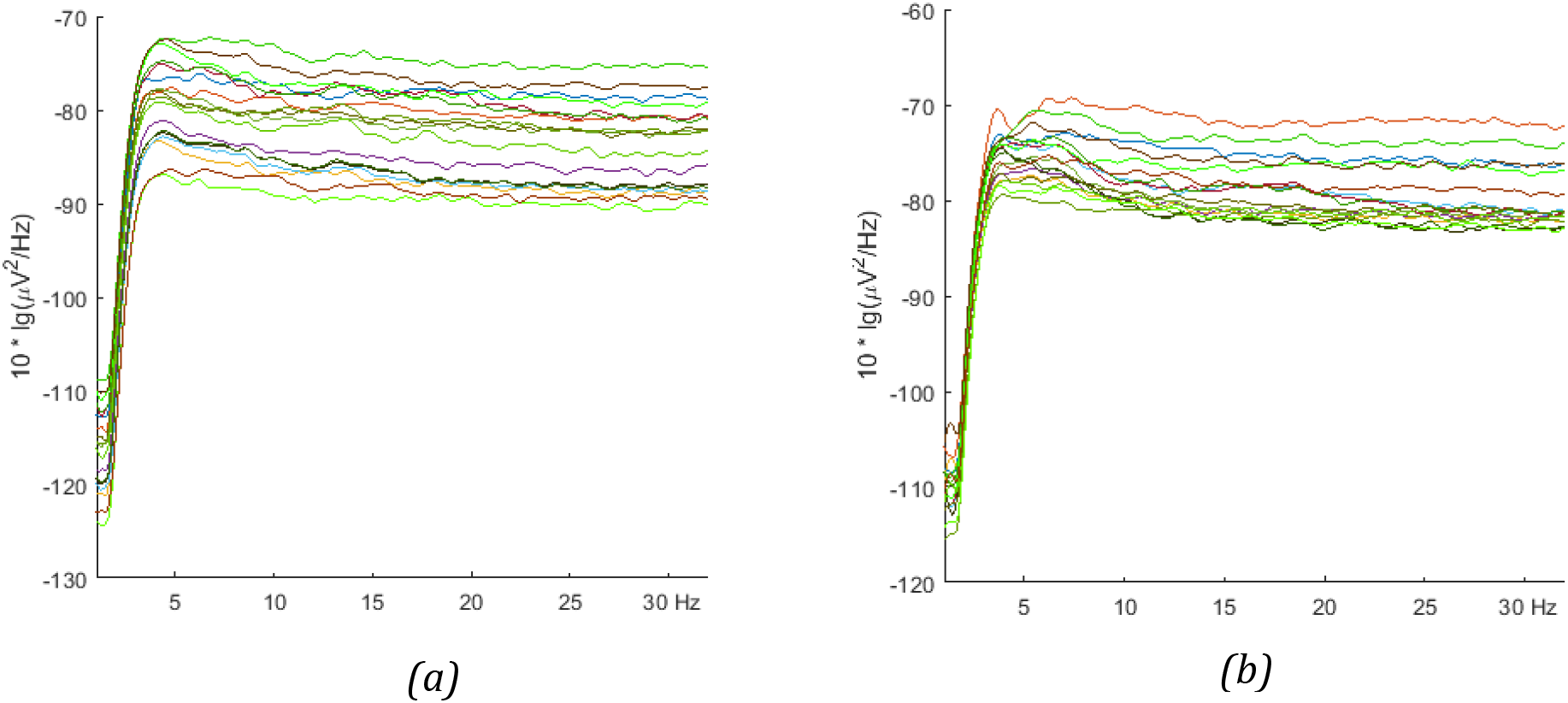
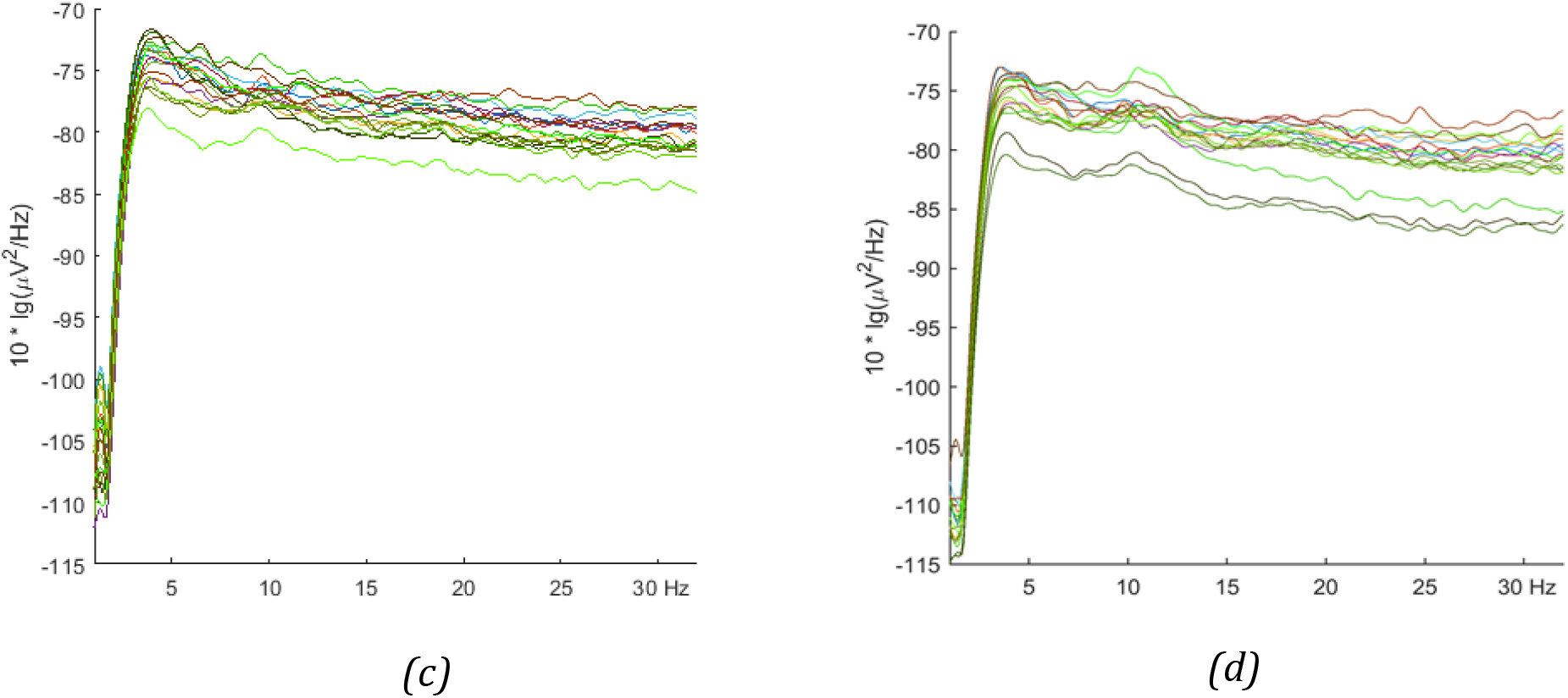
Power Spectrum for diverse visual stimuli (a) Entertainment video stimulus (b) Active Learning video stimulus (c) Infotainment video stimulus (d) Educational video stimulus

**Fig. 6:**
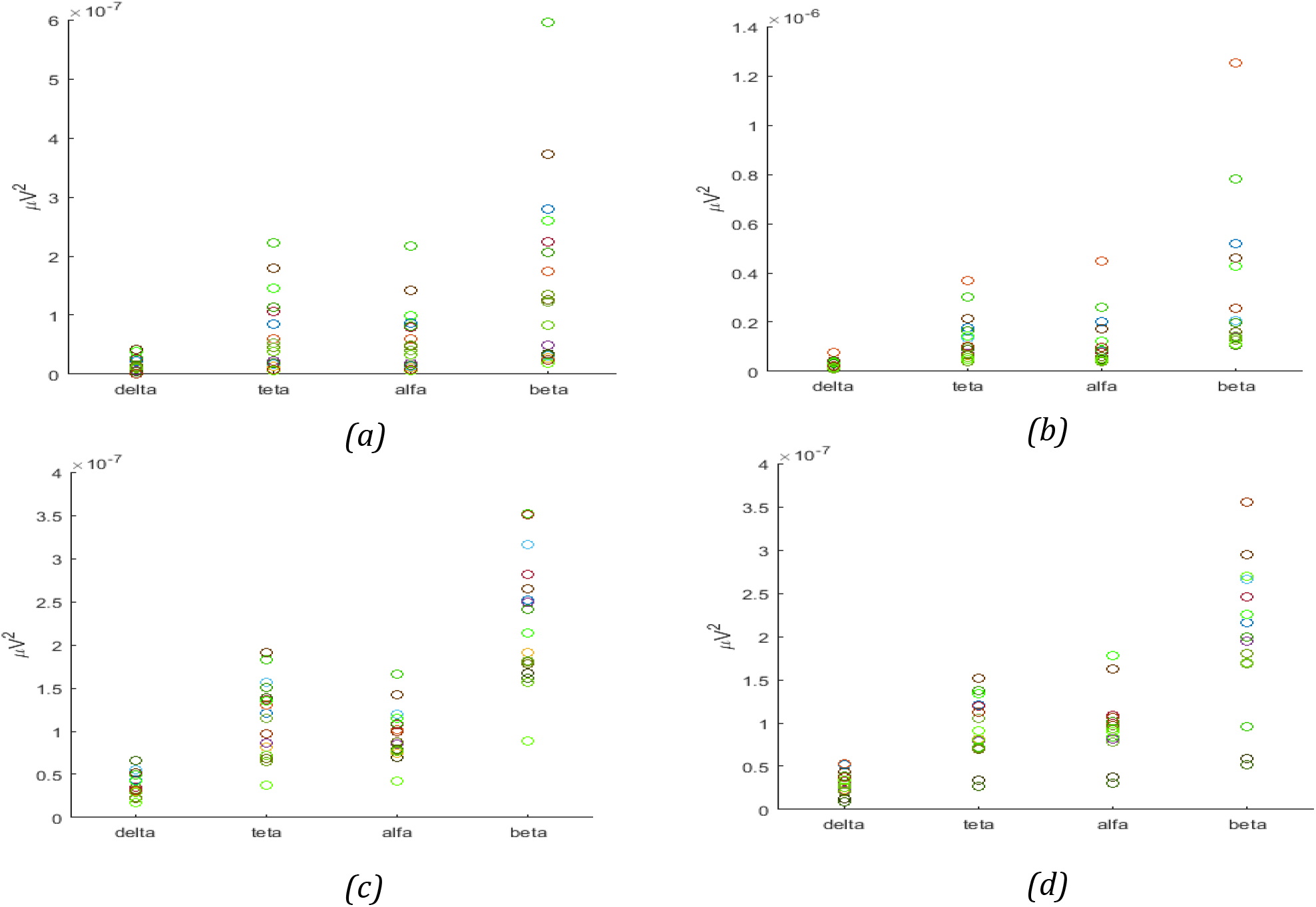
Absolute Power graph for diverse visual stimuli (a) Entertainment video stimulus (b) Active Learning video stimulus (c) Infotainment video stimulus (d) Educational video stimulus

It can be observed from Fig. 6(c) that significantly high power of delta and alpha activities are present during infotainment video stimulus. Energy contributed by various rhythms to different electrode locations is presented by the topographical brain maps in Fig. 7(a-d).

**Fig. 7:**
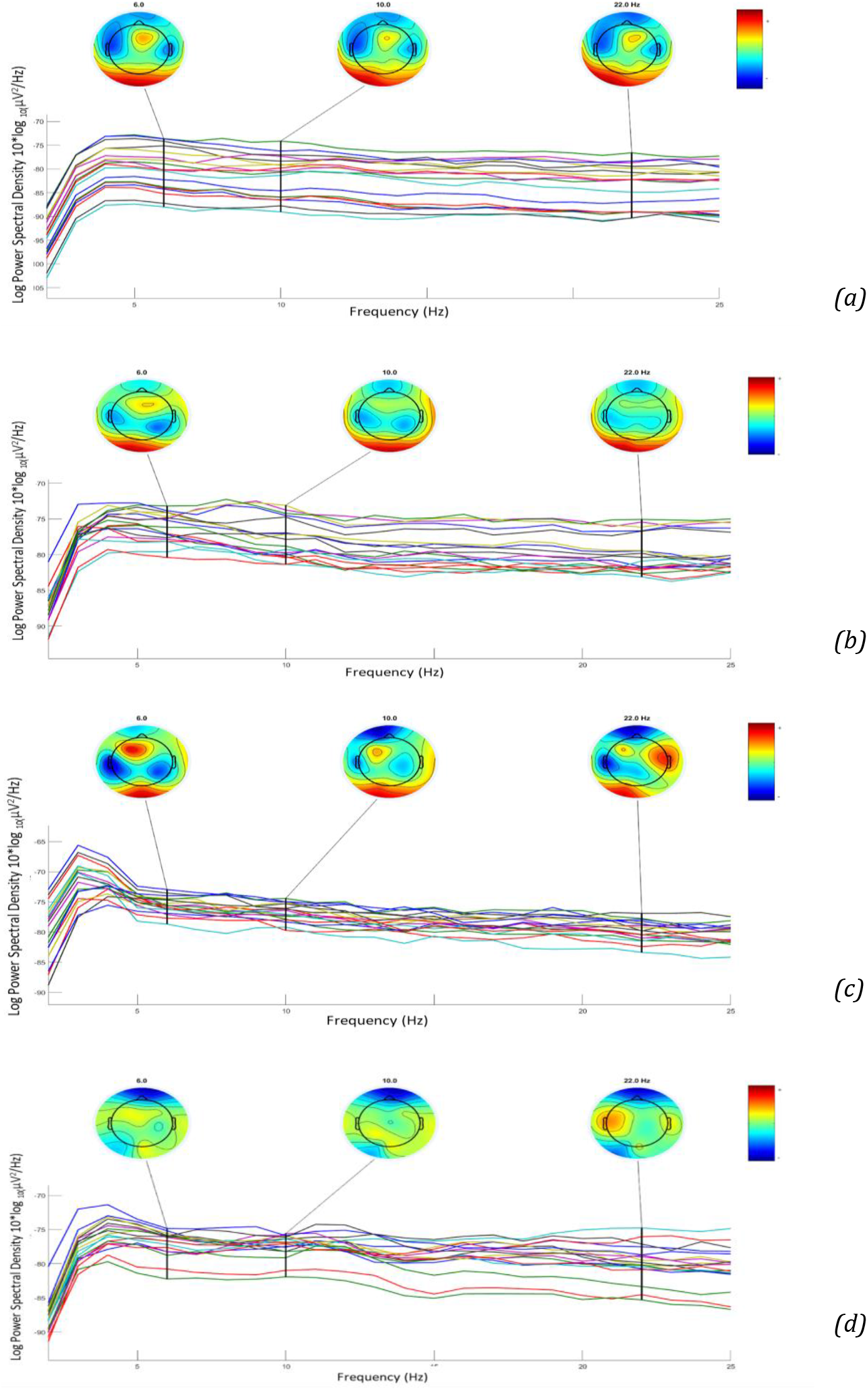
Topographical scalp-maps of the EEG activity distribution (a) Entertainment video stimulus (b) Active Learning video stimulus (c) Infotainment video stimulus (d) Educational video stimulus

It is observed from Fig. 8(a-d) that significantly elevated power levels of beta waves in the occipital and parietal regions of the brain when subjected to Active Learning. Similar observations are made when the students are exposed to entertainment video content. Fairly low power is observed from the parietal region of the brain during educational video content. From the given analysis, more engagement of the human brain is evident during active learning and entertaining video content. This concludes the hypotheses that active learning is the best form of leaning. In addition, the educational video content, if not designed with care, fails to grab attention of students

## Conclusion and Future Scope

In present work, the EEG activity of the university students was recorded, visualized and analyzed, when the students were exposed to a variety of visual stimuli such as entertainment, active Learning, infotainment and educational video. The recorded EEG activity was pre-processed to remove artifact and noise following a predefined pre-processing pipeline. Adjust method was used to remove artifactual ICs and reconstruct clean EEG activity. The visual inspection of the recorded activity is carried out using Power Spectral Density plots, Absolute Power graphs and topographical maps. Different plots and graphical methods are used to get great insight into on-going EEG activity during different experimental situations. In our analysis it is observed that significantly high EEG activity is present in the higher EEG frequency band viz. Beta band during entertainment and active learning experiments. Similar observations are drawn from Absolute Power graph and topographical maps. On contrary, the significant power drop is noticeable from lower to higher frequency bands in infotainment and educational video experiments. A clear distinction can be made among two set of EEG activity pertaining to entertainment, active Learning, and infotainment, educational video stimuli. Present study confirms high level of alertness during entertainment and active learning video stimuli. The study may help in designing suitable educational video contents to gain better attention and alertness levels in the students. This study signifies the importance of active learning in learning experience. Psychiatrists can use this information to analyse the brain activity of their patients. It can also be used to assess self academic performance, as well as the neurophysiological mechanisms underlying basic creative behaviours like idea generation and evaluation.

## Acknowledgements

The authors would like to express their gratitude to the Thapar Institute of Engineering & Technology, Patiala, Punjab for providing infrastructure and support to carry out this project work. The project was financially supported by seed money grant “Analysis of Electroencephalogram Signals for implementation of P300-based Brain Computer Interface” of the institute.

